# High-throughput mapping of meiotic crossover and chromosome mis-segregation events in interspecific hybrid mice

**DOI:** 10.1101/338053

**Authors:** Y. Yin, Y. Jiang, J. B. Berletch, C. M. Disteche, W. S. Noble, F. J. Steemers, A. C. Adey, J. A. Shendure

## Abstract

We developed “sci-LIANTI”, a high-throughput, high-coverage single-cell DNA sequencing method that combines single-cell combinatorial indexing (“sci”) and linear amplification via transposon insertion (“LIANTI”). To characterize rare chromosome mis-segregation events in male meiosis and their relationship to the landscape of meiotic crossovers, we applied sci-LIANTI to profile the genomes of 6,928 sperm and sperm precursors from infertile, interspecific F1 male mice. From 1,663 haploid and 292 diploid cells, we mapped 24,672 crossover events and identified genomic and epigenomic contexts that influence crossover hotness. Surprisingly, we observed frequent mitotic chromosome segregation during meiosis. Moreover, segregation during meiosis in individual cells was highly biased towards either mitotic or meiotic events. We anticipate that sci-LIANTI can be applied to fully characterize various recombination landscapes, as well as to other fields requiring high-throughput, high-coverage single-cell genome sequencing.

**One Sentence Summary:** Single-cell genome sequencing maps crossover and non-meiotic chromosome segregation during spermatogenesis in interspecific hybrid mice.

## Main Text

In normal mitotic cell divisions, diploid chromosomes undergo replication to generate four copies of DNA, and sister chromatids segregate apart into reciprocal daughter cells. Daughter cells receive one copy of each maternally and paternally inherited DNA sequence and almost always maintain heterozygosity at the centromere-proximal sequences (**Fig. S1A**). Rarely, chromosomes undergo mitotic crossover between chromosome homologs, which can sometimes result in diploid cells with loss-of-heterozygosity (LOH) at sequences centromere-distal to the crossover if the two recombined chromatids segregate into different daughter cells (**Fig. S1B-C**).

In meiosis, sister chromatids first co-segregate into the same daughter cell, and homologs segregate into reciprocal daughter cells in the Meiosis I (“MI”) stage, also known as meiotic or reductive chromosome segregation, resulting in diploid cells with loss-of-heterozygosity (LOH) at the centromere-proximal sequences (**Fig. S1D-E**). For the successful reductive segregation of chromosomes in MI (**Fig. S1D**), crossovers initiated by Spo11-catalyzed DSBs (*1*) provide the covalent link and necessary tension (*2*) between chromosome homologs. Rarely, chromosomes will segregate in a meiotic fashion without any inter-homolog crossover, resulting in uniparental disomy (UPD). These diploid cells then undergo mitosis-like chromosome segregation, such that sister chromatids segregate apart to form haploid gametes (**Fig. S1E**).

To date, most work on the relationship between crossover position and chromosome segregation has been performed by imaging (*3*), which fails to fully characterize the underlying genomic sequences that are prone to meiotic crossover by physically linking homologs. Genome-wide meiotic crossover maps have been generated by mapping tetrads in yeast (*4, 5*), single human sperm and complete human female meioses (*6–9*). With the exception of the studies of human female meiosis, which altogether analyzed 87 complete meioses (*6, 7*), most crossover maps are limited in three respects: 1) mature haploid gametes are analyzed where the cells have completed both rounds of meiotic division, which prevents direct observation of the more informative intermediate diploids to evaluate whether and how often chromosomes undergo meiotic or mitotic segregation (**Fig. S1**); 2) abnormal cells are selected against due to their failure to proceed to the mature gametic state; 3) analyses are limited in throughput and to a few hundred cells at the most, and as such miss out on rare events.

In principle, high-throughput, high-coverage single-cell DNA sequencing could be applied to characterize meiotic crossover and chromosome segregation profiles in an unbiased fashion. However, current single-cell DNA sequencing technologies have two major limitations. First, most methods require isolating individual cells and are consequently low-throughput. Second, most amplification methods are PCR-based and thus suffer from exponential amplification bias. To resolve the first issue, we and colleagues developed single-cell combinatorial indexing (‘sci-’), wherein one performs several rounds of split-pool molecular barcoding to uniquely tag the nucleic acid contents of an exponential number of single cells. Sci-methods have been successfully developed for profiling single-cell chromatin accessibility (sci-ATAC-seq), transcriptomes (sci-RNA-seq), genomes (sci-DNA-seq), methylomes (sci-MET), and chromosome conformation (sci-Hi-C) (*10–14*). To resolve the second issue, linear amplification via T7-based transcription provides a potential solution (*15–17*). Recently, Chen *et al*. developed Linear Amplification via Transposon Insertion (“LIANTI”), which uses Tn5 transposon to fragment the genome and simultaneously insert a T7 RNA promoter for *in vitro* transcription (IVT). RNA copies generated from the DNA template cannot serve as template for further amplification; therefore, all copies derive directly from the original DNA template. By avoiding exponential amplification, LIANTI maintains uniformity and minimizes sequence errors. However, the method is low-throughput because it requires serial library preparation from each single cell (*18*).

Here we describe sci-LIANTI, which integrates three-level single-cell combinatorial indexing and LIANTI. The sci-LIANTI method improves the throughput of LIANTI to at least thousands of cells per experiment while retaining its advantage in linear amplification. We apply sci-LIANTI to map meiotic crossover and chromosome mis-segregation events in a large number of premature and mature male germ cells from an interspecific F1 mouse cross between *Mus musculus* and *Mus spretus*.

### Overview of sci-LIANTI

The three-level combinatorial indexing and amplification schemes of sci-LIANTI are shown in **Fig. 1A**: (i) Cells are fixed with formaldehyde and nucleosomes are depleted by SDS (*12*). The resulting nuclei are then evenly distributed to 24 wells. (ii) A first round of barcodes are added by indexed Tn5 insertion within each of the 24 wells. Unlike LIANTI, wherein the Tn5 transposase contains a T7 promoter without a barcode, a spacer sequence is included 5’ to the barcodes, which serves as a “landing pad” for the subsequent ligation step (see **Fig. S2** and **Supplemental Text** for details of Tn5 transposase design). (iii) All of the nuclei are pooled and redistributed evenly into 64 new wells; a second round of barcodes is added by ligation, which includes a T7 promoter sequence positioned outside of both barcodes. (iv) All of the nuclei are once again pooled together and sorted by fluorescence-activated cell sorting (FACS) cytometry and distributed to a final round of wells at up to 300 cells per well. Note that nuclei of different ploidies can be gated and enriched by DAPI (4’,6-diamidino-2-phenylindole) staining. Also, simple dilution is an alternative to FACS that can reduce the loss rate. (v) Sorted nuclei are lysed and subjected to *in situ* gap extension to form duplex T7 promoter. This is followed by IVT, reverse transcription (RT), and second-strand synthesis (SSS) to amplify genomes in a linear fashion. A third round of barcodes is added during the SSS step, along with unique molecular identifiers (UMIs) to tag individual IVT transcripts. (vi) Duplex DNA molecules (**Fig. 1B**), each containing three barcodes that define their cell of origin are compatible with conventional library construction methods (*i.e*. appending sequence adaptors by ligation or tagmentation).

**Fig. 1.**
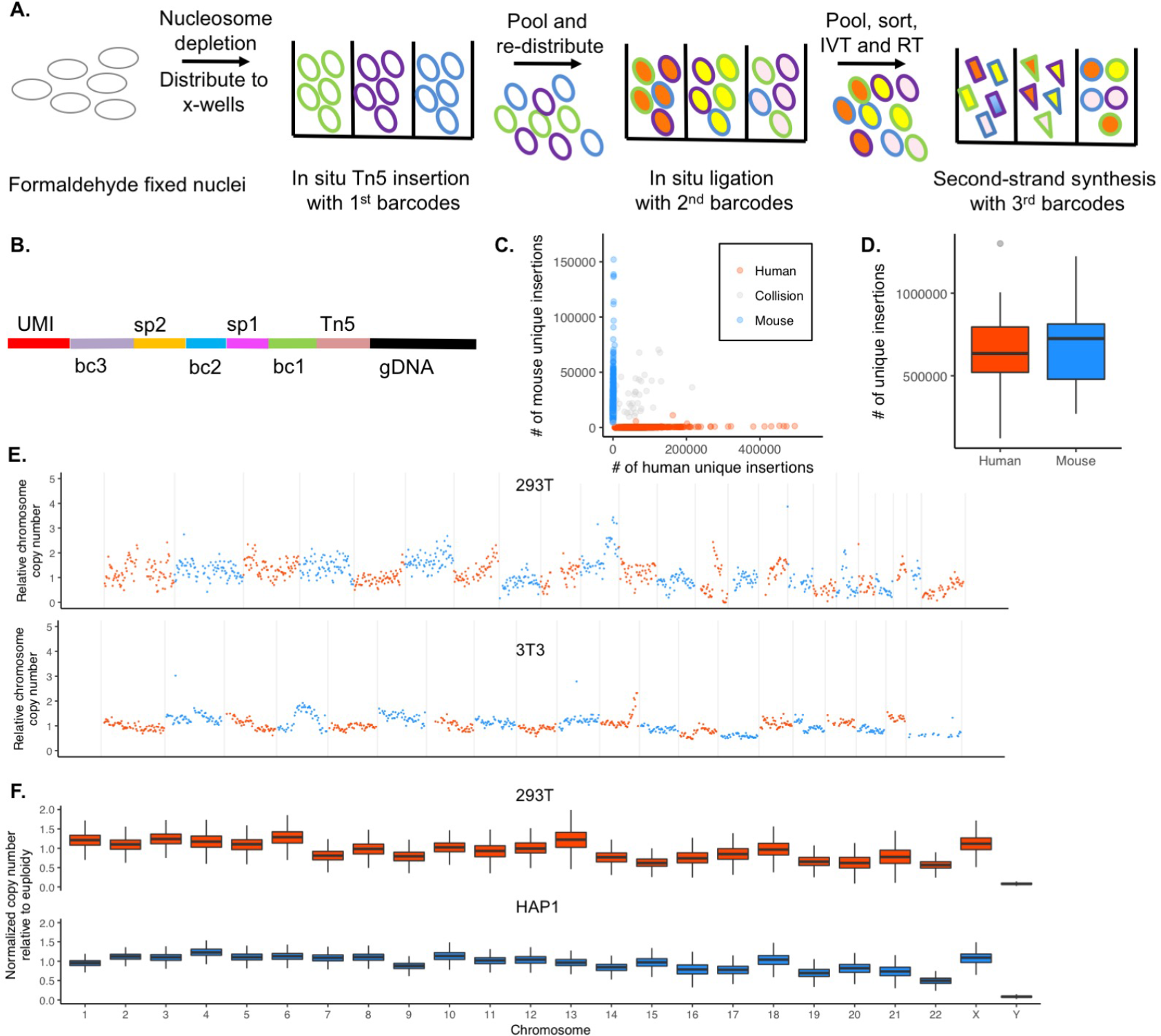
sci-LIANTI enables high-throughput, single cell, linear whole genome amplification. **(A)** Schematic of the sci-LIANTI workflow with three levels of indexing. **(B)** Barcode structure of the resulting amplified DNA duplex that is compatible with various library preparation methods. bc, barcode; sp, spacer; gDNA, genomic DNA. **(C)** Scatter plot of numbers of unique Tn5 insertion sites from human and mouse cells at low sequencing depth, 24 bc1 × 64 bc2 × 6 bc3 sci-LIANTI, 100 to 300 cells sorted per well. Blue, inferred mouse cells (percentage of mouse reads >90%, n=316); red, inferred human cells (percentage of human reads >90%, n=721); grey, collisions (n=45, 4%). **(D)** Box plots showing the number of unique Tn5 insertion sites per cell at an average of 2.4 M raw reads per cell and 1.78x depth. Depth is defined as the ratio between the number of unique transcripts and the number of unique Tn5 insertion sites. Thick horizontal lines, medians; upper and lower box edges, first and third quartiles, respectively; whiskers, 1.5 times the interquartile range; circles, outliers). See also **Fig. S3** and **Supplemental Text** for characterization of libraries made with improved versions of the protocol. **(E)** Example chromosome CNV plots. Upper, HEK293T cell, 2.6 M raw reads, 2.4 M unique molecules, 1.3 M unique Tn5 insertion sites with MAPQ > 1. Lower, 3T3 cell, 2.7 M raw reads, 2.4 M unique molecules, 1.2 M unique Tn5 insertion sites with MAPQ > 1. **(F)** Box plots for copy number variation aggregating 822 293T cells or 1,453 HAP1 cells. Y-axis depicts reads fraction per chromosome normalized by chromosome length such that euploid chromosome without segmental copy gain or loss is expected to have a value of 1.

As a proof-of-concept, we mixed mouse and human cells and performed sci-LIANTI. For over 95% of the resulting single-cell genomes, the vast majority of reads map either to the mouse or human genome; occasional ‘collisions’ result from chance use of the same combination of barcodes by two or more cells (**Fig. 1C**). The performance of sci-LIANTI is compared to LIANTI as well as our previous PCR-based sci-DNA-seq method in **Table S1**. We highlight several advantages of sci-LIANTI: 1) We generally recover 90% of sorted cells as compared to 60% recovery with PCR-based sci-DNA-seq (*12*); 2) With 40% fewer raw reads (329M by sci-LIANTI vs. 549M by sci-DNA-seq), sci-LIANTI produced sequence coverage at ~97,000 unique Tn5 insertions per cell, as compared to ~30,000 unique insertions by sci-DNA-seq, a >3-fold improvement. Sequencing a smaller number of cells to a higher depth, we observed ~660,000 unique Tn5 insertions per cell while maintaining higher library complexity than sci-DNA-seq, suggesting a further improvement of >20-fold; 3) The rate of mappable reads is improved from ~40% with LIANTI to ~80% with sci-LIANTI. This is likely because LIANTI is entirely in-tube and therefore it is hard to remove artifactual sequences, *e.g*. secondary to self-insertion of Tn5, whereas with sci-LIANTI, nuclei are pelleted several times to remove excess free DNA; 4) Unlike PCR-based amplification where duplicate reads are not informative for SNP calling, sci-LIANTI’s ‘duplicate’ reads almost always result from independent IVT transcripts polymerized from the original template, and are therefore useful for *de novo* SNV discovery or for genotyping known SNPs.

With sci-LIANTI, Tn5 inserts on average every 0.5-1.5 kb of the human genome per cell, and IVT amplifies ~1,000-fold. This corresponds to 2 to 6 million unique Tn5 insertions, and 2 to 6 billion unique genome-derived IVT transcripts, per single cell. It is obviously currently impractical to sequence the resulting library to saturation with respect to the number of unique IVT transcripts. We define the ratio between the number of unique transcripts sequenced to the number of unique Tn5 insertions sites mapped as the ‘depth of sequencing’ for each library. In this study, most of the libraries are sequenced at a depth of 1.1x to 2x, resulting in 0.5% to 5% coverage of the genome of each cell. The distribution of unique Tn5 insertion sites per cell in the human/mouse proof-of-concept experiment is shown in **Fig. 1D**, and for other experiments in **Fig. S3**. The estimated relative chromosomal copy numbers for representative single cells is shown in **Fig. 1E**, and their distributions across all cells in **Fig. 1F**. To extrapolate expected genome coverage per single cell at higher sequencing depth, we fit the number of unique insertion sites as a function of sequencing depth (**Fig. S3G**). We expect to observe 4.2M and 6.0M unique insertions per cell at a sequencing depth of 5x and 10x, respectively, which corresponds to 16% and 22% coverage of the genome of individual cells.

Note that at the molecular level, we have modified both the “sci” and “LIANTI” methods in several ways. In brief, we changed design of the Tn5 transposon to be compatible with ligation, added a loop structure of the T7 promoter to facilitate intra-molecular ligation, and changed the RT scheme such that we only require successful ligation at one of the two ends of the first-round barcoded molecules. Supposing that a single ligation event has 50% efficiency, this modification renders a 75% success rate at the ligation step instead of 25% (comparison shown in **Fig. S3** and details in **Supplemental Text**). We depict the structures of the molecules after each barcoding step in **Fig. S2** and discuss rationales for these designs in the **Supplemental Text**. Scalability and cost are also discussed in the **Supplemental Text** and **Table S2**. For libraries of 100, 1000 or 10,000 single cells, we estimate the cost of sci-LIANTI to be 14%, 1.5% and 0.26% of that of LIANTI. We also show that using the three rounds of barcoding substantially reduces costs when compared to a two round strategy.

### Single-cell DNA profiling of mouse germ cells

We next applied sci-LIANTI to study mouse meiosis in an interspecific mouse cross. Unlike inbred mice whose epididymides store millions of mature sperm, the epididymides of F1 males (*19*) resulting from crossing female *Mus musculus* C57BL/6 (subsequently ‘B6’) and male *Mus spretus* SPRET/Ei (subsequently ‘Spret’) contain extremely low numbers of morphologically mature sperm and limited numbers of round germ cells of unknown ploidy (**Fig. S4A-B**). We pooled cells isolated from 6 and 3 epididymides from (B6 x Spret) hybrid males aged 70 days and 88 days, respectively, in two separate experiments, and fixed with 1% formaldehyde. For each experiment, after nucleosome depletion, we distributed 30,000 nuclei per well and performed *in situ* indexed Tn5 insertion across 24 wells to add the first-round barcodes. We then pooled all ~700k nuclei and redistributed these to 64 wells to add the second-round barcodes and T7 promoter by ligation. After again pooling all cells, we split the cell mixture 1:6, FACS-sorted the majority of cells (6/7), and diluted the rest (1/7). The resulting wells contained 100 to 360 cells per well (predicted barcode collision rate of 4% to 11%). Interestingly, we observed a much higher fraction of 2C (DNA content of an unreplicated diploid cell) cells during FACS (**Fig. S4C-D**) than would be expected for a ‘normal’ epididymidis, which is dominated by 1C sperm. The number of cells recovered and their estimated cell ploidy is listed in **Table S3**. We proceeded with linear amplification, second strand synthesis to add the third-round barcode, library preparation, and sequencing.

### M2 diploid cells have clustered mitotic or meiotic chromosome segregation

From these two experiments, we profiled the genomes of 2,689 (92% of 2,919 cells sorted with >10k raw reads) and 4,239 (94% of 4,497 cells sorted with >30k raw reads) single cells. The number of uniquely mapping reads are shown in **Fig. S3D**. At a sequencing depth of 1.6x and 1.4x for the two libraries (details in **Fig. S3**), we obtained a median of ~70k and ~144k unique Tn5 sites per cell, corresponding to 0.7% and 1.4% median genome coverage.

To identify crossover breakpoints, we implemented a hidden Markov model (HMM) that relied on high-quality reads that could clearly be assigned to B6 or Spret (see **Supplemental Text, Table S6, Table S8**). We characterized crossovers in 1,663 haploid cells, a representative example of which is shown in **Fig. 2A**. In addition, we searched ~5,200 diploid or 4C cells for crossover events. To our surprise, we identified 292 diploid cells with a significant number of crossovers, which we term “M2 diploids” (**Fig. 2B** and 2C). Even more surprisingly, a substantial proportion of these cells exhibited *mitotic*, rather than meiotic, segregation.

**Fig. 2.**
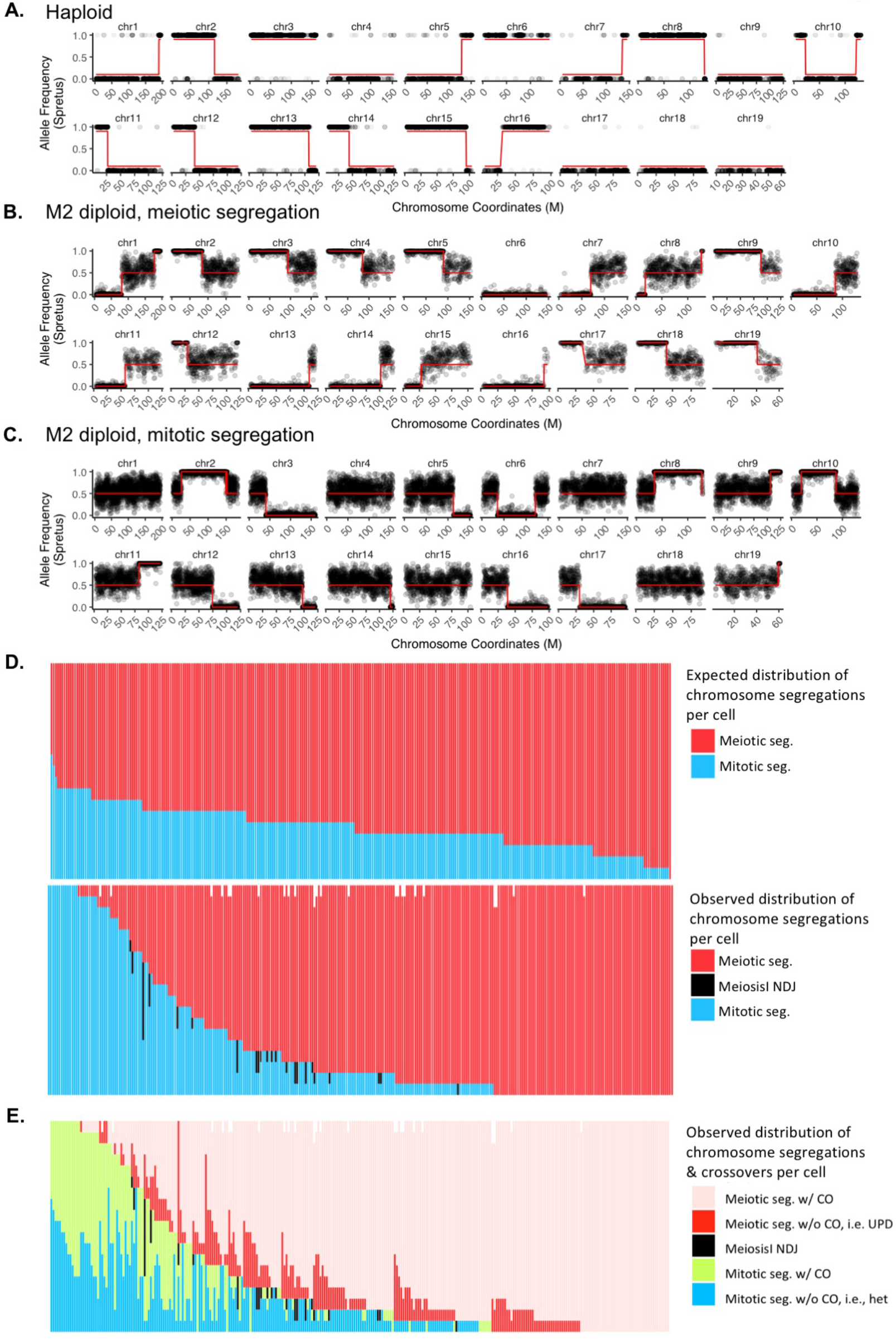
sci-LIANTI of the hybrid mouse male germline reveals numerous examples of nonindependent mitotic segregation. In (A), (B) and (C), red line depicts fitted crossover transition via HMM. Centromere is located at the leftmost for picture of each chromosome. **(A)** Example crossover plot for haploid. Grey dot has a value of 1 for Spret allele and 0 for B6 allele. In (B) and (C), grey dot shows allele frequency of Spret averaging 40 SNP sites. **(B)** Example LOH plot for M2 diploid with meiotic segregation (see also **Fig. S1D**). LOH is present at the centromere-proximal region of the crossover sites. **(C)** Example LOH plot for M2 diploid with mitotic segregation (see also **Fig. S1B**). LOH is present at the centromere-distal region of the crossover sites, unlike in (B). **(D)** Number of meiotic **(red)** and mitotic (blue) chromosomes for each of the M2 diploid cell. Each column represents one single M2 diploid cell (19 chromosomes per cell, distributed as indicated by colors). **Top**: expected distribution of meiotic vs. mitotic segregation based on the binomial distribution and assuming the probability of meiotic segregation p equals 0.76, the MLE from the observed data. **Bottom**: observed data in M2 diploid cells. In rare cases (27/5,548 chromosomes), we were not able to distinguish meiotic vs. mitotic segregation due to sparse SNP coverage (white space at the top of the panel). Black bar depicts Meiosis I nondisjunction (NDJ, 40 chromosomes in total) where we observed 0 or 4 copies of the chromatids. **(E)** Breakdown of (D) by the number of chromosomes with or without crossovers (abbreviated as “CO”). Cells are sorted first by the number of meiotically segregated chromosomes (pink and red, in ascending order) and then by the number of observed mitotic crossovers (light green, in descending order). As expected, none of the Meiosis I NDJ chromosomes contain crossovers.

After a crossover occurs between two chromosome homologs, if the chromosome segregates in a meiotic fashion, the region between the centromere and the position of crossover will become homozygous, whereas heterozygosity will be maintained downstream of the crossover (**Fig. S1A**). However, if the chromosome segregates in a mitotic fashion, LOH is observed centromere-distal to the crossover if the recombined chromatids segregate apart (**Fig. S1B**). We show one example of an M2 diploid whose chromosomes undergo meiotic segregation in **Fig. 2B**, and one example of an M2 diploid whose chromosomes unexpectedly undergo mitotic segregation in **Fig. 2C**. In total, across 292 M2 diploid cells, we observed 4,162 examples of chromosomes undergoing meiotic segregation, among which 3,740 harbor crossovers (90%), and 1,310 examples of chromosomes undergoing mitotic segregation, among which we observe 636 crossovers (49%). Of note, however, the number of mitotic crossover events may be higher, as we cannot identify a subset of crossover outcomes (**Fig. S1C**); meanwhile, we can detect all crossovers for meiotically segregated chromosomes.

Although we observe many examples of cells where some chromosomes exhibit meiotic segregation and other chromosomes exhibit mitotic segregation, the segregation pattern of individual chromosomes within M2 diploid cells does not appear to be independent. If chromosomes in each cell chose meiotic vs. mitotic segregation independently, we would expect a binomial distribution of meiotically and mitotically segregated chromosomes, centered on the maximum likelihood estimate (MLE) of the probability, p, of meiotic segregation (p=0.76 from the data, 4162/5472), with roughly three quarters of chromosomes segregating meiotically and one quarter segregate mitotically (**Fig. 2D**, top panel). However, among the 292 M2 diploid cells that we profiled, we observe 202 cells that have at least 15 chromosomes segregating in a meiotic fashion, and 38 cells that have at least 15 chromosomes that segregated in a mitotic fashion (**Fig. 2D, bottom panel**; this contrasts with 148 and 0 cells expected, respectively, under assumption of independence). That individual M2 diploid cells are biased towards overwhelmingly undergoing meiotic or mitotic segregation suggests a global sensing mechanism for deciding whether a cell proceeds with meiosis or returns to mitosis.

We can further classify cells by whether the chromosomes in M2 diploids have a crossover (**Fig. 2E**). Meiotically segregated chromosomes appear to have more crossovers (pink in **Fig. 2E**) than mitotically segregated chromosomes (green in **Fig. 2E**). However, unlike in meiotically segregated chromosomes where we can detect all the crossovers as centromeric LOH, mitotically segregated chromosomes only have LOH if the two recombined chromatids segregate apart into reciprocal daughter cells (**Fig. S1B**). If instead recombined chromatids co-segregate, heterozygosity will be maintained throughout the chromosome despite the undetectable linkage switch (**Fig. S1C**). In **Fig. 2E**, the ratio of having (shown in green) vs. not having (shown in blue) an observable LOH in mitotically segregated chromosomes is roughly one. This could either mean that mitotically segregated chromosomes have a 50% chance of segregating recombined chromatids together, if those completely heterozygous chromosomes (shown in blue) do have a linkage switch; or alternatively that mitotically segregated chromosomes always segregate recombined chromatids apart, and the crossover frequency is reduced by half compared to meiotically segregated chromosomes.

### Distribution of meiotic crossover at the chromosomal level

We next sought to investigate genomic correlates of crossover events. Altogether, we analyzed 1,663 haploid cells harboring 19,601 crossover breakpoints and 292 M2 diploids with 5,071 crossover breakpoints, to our knowledge an unprecedented dataset with respect to the combination of breadth and resolution for querying crossovers in mammalian meiosis. We first considered the distribution of meiotic crossovers across chromosomes. Crossover density is defined here as the average number of crossovers per cell per division per Mb multiplied by 2 (in haploids) or 1 (in M2 diploids). We observed a strong negative correlation between chromosome size and crossover density in haploid cells (**Fig. 3A**, r = −0.66, p = 0.002). Consistent with previous findings (*20*), this correlation is only partly explained by Spo11 oligonucleotide complex density (r = −0.46, p < 0.05), suggesting that smaller chromosomes sustain more DSBs and those DSBs are more likely to give rise to crossovers. The negative correlation is even stronger in M2 diploid cells (**Fig. 3B**, r = −0.8, p = 5e-5). In **Fig. S5A-B**, we consider instances of multiple crossovers per chromosome per cell as a single event, which strengthens the negative correlation even further (r = −0.87, p = 2e-6 for haploids and r = −0.9, p = 2e-7 for M2 diploids). These observations suggest that smaller chromosomes are hotter for crossovers, and particularly for having at least one crossover per cell division. Haploid cells had an average of 0.62 crossovers per chromosome per cell, while M2 diploids had an average of 0.91 per chromosome per cell (**Fig. 3C-D**). The crossover rate in M2 diploids is only 9% lower than crossover counts measured by Mlh1 foci in 4C spermatocytes in B6 inbred mice (*21*). The crossover rate in haploid cells is 45% lower than observed in single human sperm sequencing (*8, 9*). The latter difference could largely be due to the telocentric nature of mouse chromosomes. We note that among the 80 M2 diploids that segregated all 19 autosomes meiotically, which allows us to detect all crossovers, 41 have a crossover on every autosome despite presence of almost 2% nucleotide polymorphism between homologous chromosomes in (B6 x Spret) F1 hybrids.

**Fig. 3.**
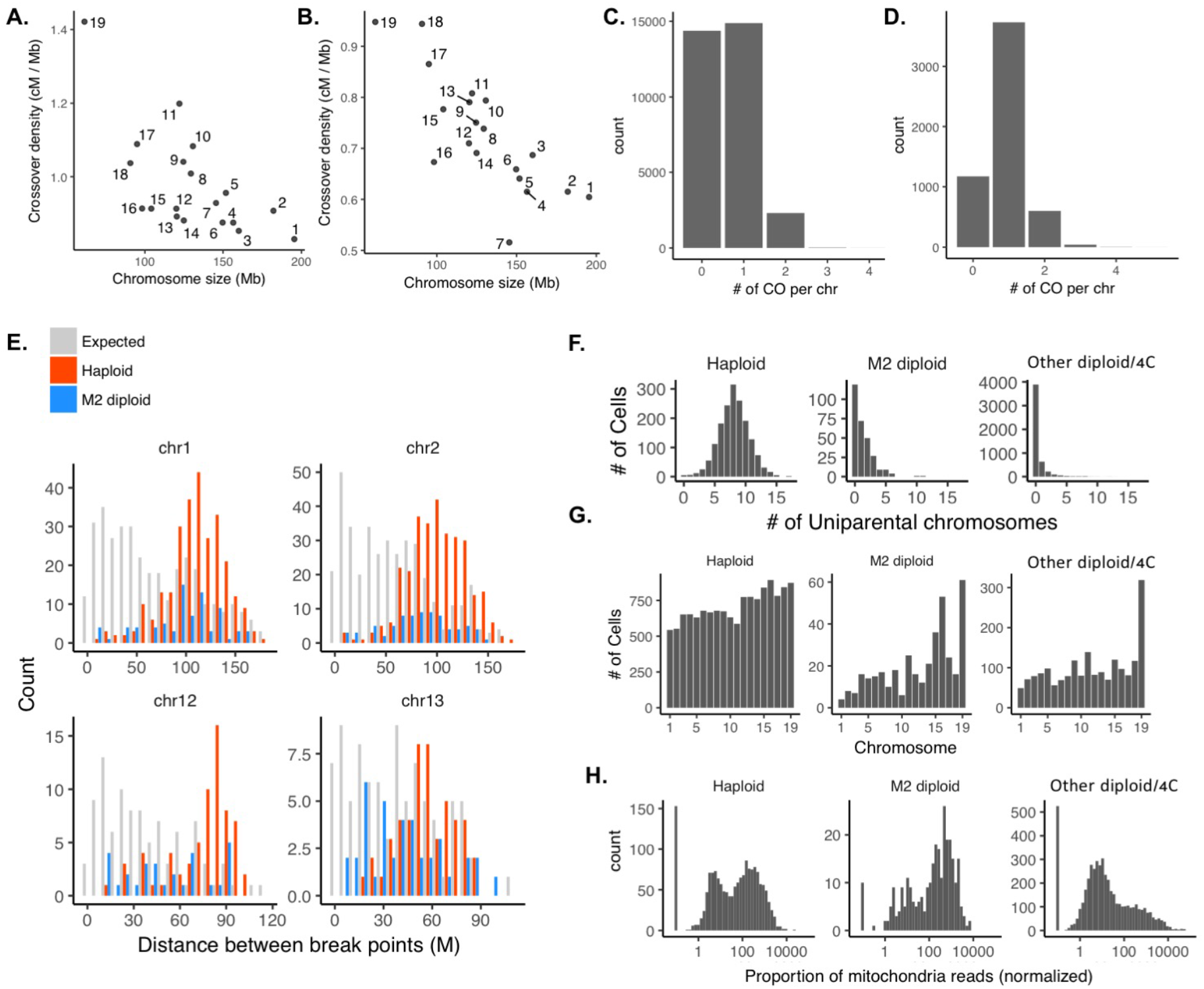
Meiotic crossover and uniparental chromosome distributions at the chromosome scale. **(A)** Number of crossovers normalized by chromosome size (cM/Mb) negatively correlates with chromosome size in haploid cells (r = −0.66, p = 0.002). (B) Same as (A) in M2 diploid cells (r = −0.79, p = 5e-5). (C) Distribution of crossover (CO) counts per chromosome per haploid cell (mean = 0.62). **(D)** Same as (C) for M2 diploids (mean = 0.91). **(E)** For chromosomes with at least two crossovers, distance (Mb) between crossovers for chromosomes 1, 2, 12, 13, See **Fig. S5C** for all chromosomes. The distribution of expected numbers is generated by randomly placing 2 or 3 crossovers per chromosome. **(F)** Uniparental chromosome numbers per haploid (median = 8, mean = 8.1), M2 diploid (median = 1, mean = 1.3), or other diploid/4C (median = 0, mean = 0.4) cells. **(G)** Uniparental chromosome distribution (correlation with chromosome size shown in parentheses), haploid (r = −0.87, p = 2e-6), M2 diploid (r = −0.75, p = 2e-4), other (r = −0.67, p = 0.002). **(H)** Number of mitochondrial reads per cell, normalized by read depth, for haploid, M2 diploid, and other diploid/4C diploid cells.

To examine crossover interference, we took chromosomes with at least two crossovers and plotted the distance between adjacent crossovers, and compared this distribution to random expectation based on simulation (**Fig. 3E, Fig. S5C**). The average observed distance between crossovers was 79 Mb, which is much larger than the expectation of 42 Mb (p=3e-243). This is consistent with the repulsion of crossovers of close proximity. We also analyzed distribution of uniparental chromosomes in each single cell (**Fig. 3F**, Table S7, Table S9) and for each chromosome (**Fig. 3G**). Although shorter chromosomes exhibit elevated crossover rates when normalized by length, the rate of uniparental chromosomes without observed crossovers still negatively correlates with chromosome size (r= −0.91, p=4.6e-8, breakdown for each class of cells in **Fig. 3G**). It is worth noting that the Patski cell line, which is a spontaneously immortalized cell line derived from female B6/Spret F1 mouse, also occasionally has UPD and mitotic LOH, although at a much reduced rate. We analyzed 1,107 single cells from Patski, among which we found an average 0.36 UPD and 0.098 segmental LOH events (**Table S10**). However, we note that these events are not necessarily independent. For example, a UPD emerging early in the passage of the cell line can be shared in a large portion of the cells derived. Nevertheless, chromosome distribution of these events is plotted in **Fig. S5D**.

Lastly, we examined the normalized proportion of reads per cell that map to the mitochondrial genome (**Fig. 3H**). Haploid cells exhibit a bimodal distribution in terms of the “copy number” of mitochondria DNA. One speculative possibility is that in mature sperm, the mitochondria are near the tail of the sperm cell and are thus physically separate from the bulk nuclei, while in 1C round spermatids, the mitochondria localize closely enough to the nuclei to be crosslinked with the nuclei. We observed a negative correlation between the mitochondrial read proportion and the number of crossovers (rho= −0.11, p=3e-6). Interestingly, although of limited number, M2 diploids that segregated at least 14 of their chromosomes either mitotically and meiotically have very different distributions of mitochondrial read proportions (**Fig. S5E**). Consistent with this, the mitochondrial read proportion positively correlated with the number of meiotically segregated chromosomes in M2 diploid cells (r = 0.21, p = 0.0003).

### Distribution of meiotic crossover at the break point level

We piled up crossovers throughout the genome to generate crossover hotness maps along each chromosome and compared this with the Spo11 oligonucleotide-complex map, which identifies meiotic double strand break hotspots at the highest resolution (*20*). Haploid (purple) and M2 diploid (green) are correlated in the crossover pileup (r = 0.65, p < 2e-16), but both deviate from the Spo11 map (r = 0.059 for haploids and M2 diploids combined, p = 4e-20, r = 0.059 and r = 0.044 for haploids and M2 diploids, respectively). Note that PRDM9 evolves to bind different motifs between diverged mouse strains, even between subspecies of mouse (*3, 22*). The erosion of PRDM9 consensus binding site results in four types of DSB hotspots defined by the Spo11 oligonucleotide-complex map, those that are conserved between B6 and Spret, termed as “symmetric” hotspots (shown in **Fig. 4A**), those that are only present in B6 or Spret, termed as “asymmetric” hotspots and those do not contain PRDM9 binding site in either species (all hotspots combined regardless of symmetry are shown in **Fig. S6**). We observe a stronger correlation for “symmetric” hotspots than all hotspots (r = 0.067 for haploids and M2 diploids combined, p = 1.8e-25; r = 0.065 and r = 0.055 for haploids and M2 diploids, respectively).

**Fig. 4.**
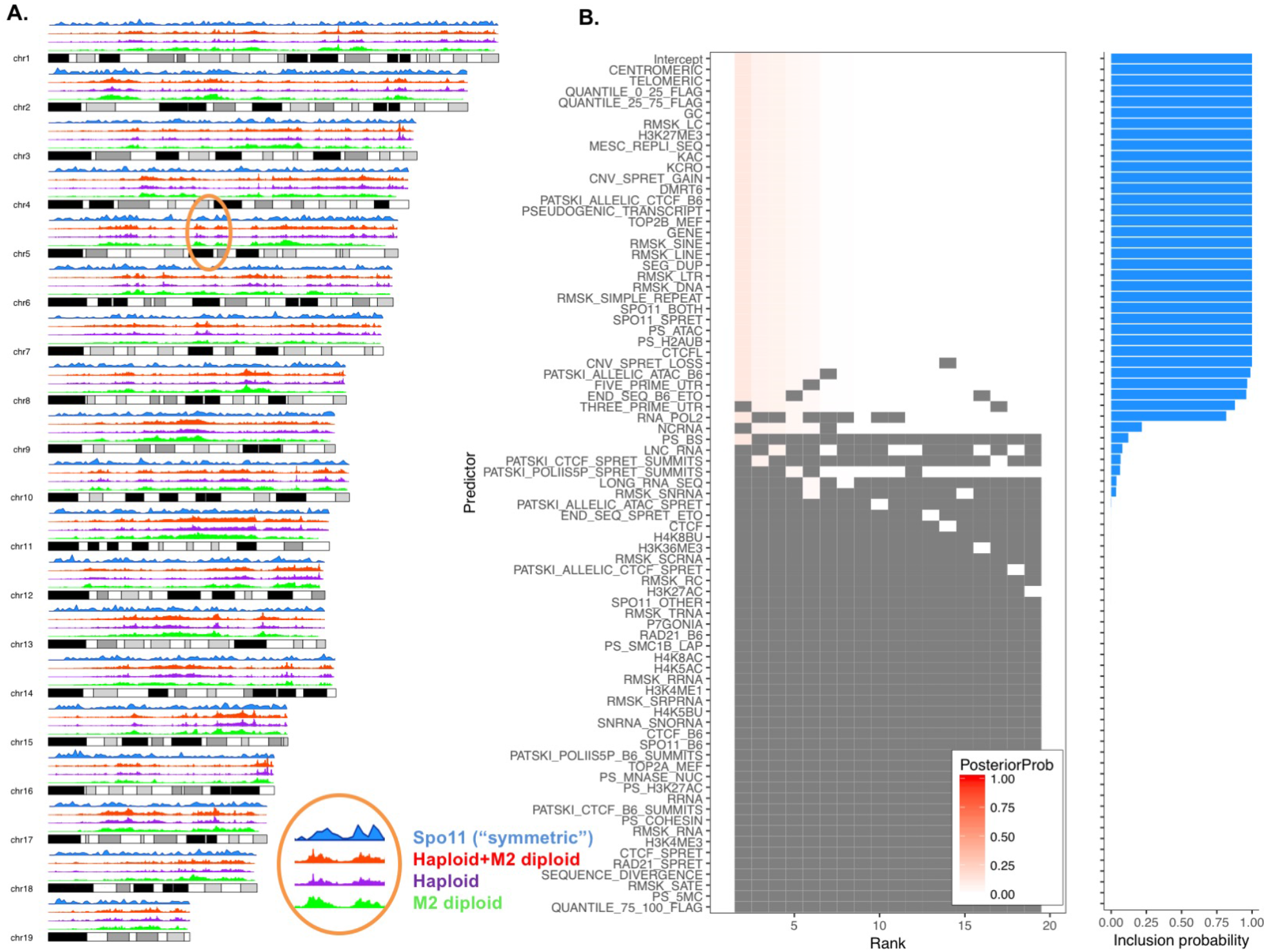
Meiotic crossover hotness and explanatory genomic features. **(A)** Crossover break point pileup profile (top to bottom: meiotic DSB hotspot by Spo11-oligo map for symmetric hotspots, *i.e.*, where PRDM9 motif is present in both B6 and Spret assembly, haploid+M2 diploid, haploid, M2 diploid). **(B)** Marginal inclusion probability for features associated with crossover hotness by BMA. The x-axis ranks models by posterior probability.

Only 10% of meiotic-specific breaks are repaired as crossovers. We next looked at what factors beyond Spo11 breaks contribute to crossover formation by building a linear model with Bayesian Model Averaging (BMA, (*23*)) and we quantified an inclusion probability for ~80 potentially explanatory variables. Features that have been known to affect meiotic crossover such as Spo11 break sites, GC content, quantiles along the chromosome, etc. are included in almost all the models with high probabilities (**Fig. 4B**), and we also found a few more features that have not previously been implicated in meiotic crossovers, such as specific families of repeats and chromatin marks. A correlation matrix between crossover hotness and all the features is plotted in **Fig. S7** and summaries of the linear model are included in **Table S4** and **Table S5**.

Although haploids and M2 diploids appear similar in the crossover pileups (**Fig. 4A** and **Fig. S6**), we asked whether there is structure to the features that influence crossover distributions across subsets of single cells. The median crossover break point resolution is ~150 kb (**Fig. 5A**). By aggregating information in each single cell, we can cluster cells based on three sets of features. For the first set of features, we summarize the number of crossover or whole-chromosome LOH for each chromosome. For the second set of features, we obtain median values for the crossover breakpoints in each single cell in terms of GC content, sequence divergence, intensity of chromatin marks, etc. For the third set of features, we sum up the counts of genome elements such as genes bodies, long terminal repeats (LTR), LINE elements included in all the breakpoints in each single cell and normalize by total tract lengths.

**Fig. 5.**
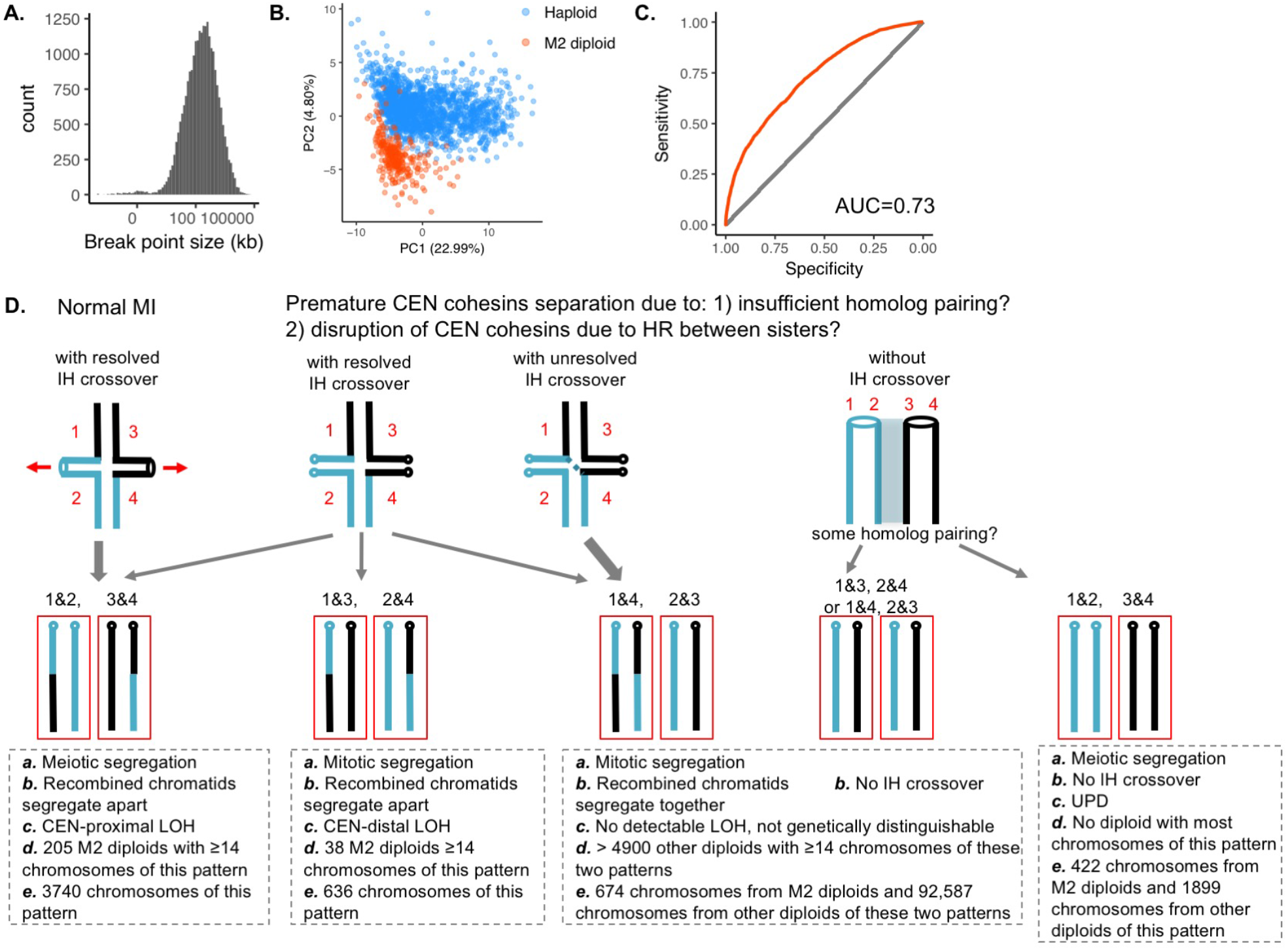
Cell clustering, predictions of meiotic crossover tracts and model for relationship between meiotic crossover and chromosome mis-segregation. **(A)** Distribution of sizes for breakpoint resolution (log normal distribution). **(B)** PCA with ~120 features shows distinct clusters corresponding to haploid and M2 diploid cells. **(C)** AUC of 0.73 quantifies accuracy in predicting if a region drawn from the mouse genome comes from crossover tracts or random tracts. We observed the AUC to remain the same whether the number of crossover and random tracts used in the ROC analysis are 1:1 or 1:9. **(D)** Model for meiotic crossover and chromosome mis-segregation. “MI”, meiosis I, “CEN”, centromere, “IH”, inter-homolog. Frequencies of patterns of segregation are indicated by the relative widths of the grey arrows.

We then used principal component analysis (PCA) on a matrix with each row as one single cell and each column as one summarized value for one feature. The first two PCs capture 27% of the total variance, and the haploids and M2 diploids are separated into two clusters (**Fig. 5B**). In **Fig. S8** we plot each feature projected on the first two PCs, and we observed that chromosome distribution of crossovers and UPDs rather than numbers of such events seem to drive the separation of haploids and M2 diploids. Crossovers in M2 diploids tend to occur in the middle of the chromosome arms, while those in haploids tend occur towards the telomeres (**Fig. S9**).

Finally, we contrasted crossover breakpoints with random regions drawn from the mouse genome. We built a linear model of binary response with 1 being crossover tracts and 0 being a random tract sampled from the genome from the same tract length distribution (details in **Supplemental Text**). Using the same 80 features as in the BMA analyses, we can predict crossover tracts with an average accuracy of 0.73 (**Fig. 5C**) for held-out data with all the features and with an average accuracy of 0.73 and 0.713 with variables of high inclusion probability (>0.5) identified by BMA and variables that are significant from simple multivariate linear regression, respectively.

## Discussion

We have developed sci-LIANTI, which provides high-throughput and linear amplification of tens-of-thousands of single-cell genomes in a two-day experiment, at a cost of $0.14 per cell for library construction. Given its high-throughput nature, sci-LIANTI enables the identification of rare and unexpected cell types in an unbiased manner, which is difficult to achieve by ‘in-tube’ LIANTI (*18*), and improves on the number of unique molecules present in each single cell from low thousands (*24*) or low tens of thousands (*12*) to hundreds-of-thousands at an affordable sequencing depth. In this study of mouse meiosis, sci-LIANTI identified an unexpected population of M2 diploid cells. The single-] cell nature of the data also allowed us to simultaneously examine meiotic crossover and chromosome mis-segregation. The improved genome coverage enabled high-resolution mapping of crossover break points, and allowed us to better characterize genomic and epigenomic features associated with crossover hotness. Although the mapping of mouse meiosis requires whole genome sequencing, we note that sci-LIANTI can be easily adapted to targeted sequencing by a small modification of the protocol (**Supplemental Text**).

Chromosomes that segregate mitotically have previously been observed in the complete analyses of human female meiosis, and termed “reverse segregation”. Acrocentric chromosomes seem to have a slightly elevated rate of reverse segregation, although the study observed only 23 meioses (*7*). Among the 292 M2 diploid cells we analyzed here, individual cells were biased towards mitotic or meiotic chromosome segregation, consistent with a global sensing mechanism for deciding whether a cell proceeds with meiosis or returns to mitotic segregation of its chromosomes. Alternatively, this could be explained by the telocentric nature of mouse chromosomes, although it is not clear why telocentric chromosomes would tend to segregate mitotically, particularly when there is a normal crossover. Our observation also raises a few interesting questions: 1) mitotic segregation in MI indicates premature centromeric cohesin separation. As to what causes this, is it possible that due to insufficient homolog pairing between B6 and Spret chromosomes, DSBs that should have been repaired off of the homolog are instead frequently repaired using sister chromatids as template? This could cause disruption of cohesins (*25*). 2) Is having one crossover between homologs sufficient to engage them, leading to meiotic segregation in MI? Based on current models, one inter-homolog crossover and sister chromatid cohesion seem to be the only prerequisites for forming chiasmata. By this model, despite insufficient homolog pairing, once a crossover is successfully formed, chromosome segregation should not be impaired. However, the large numbers of mitotically segregated chromosomes observed here, in spite of normal crossovers, could indicate that defects in initial homolog pairing impact the ultimate outcome. 3) What are the consequences of these mitotically segregated chromosomes? Do they go back to mitosis or do they proceed to Meiosis II (“MII”)? In yeast, a phenomenon called “return-to-growth” has been characterized wherein cells that initiate the meiosis program can revert to normal mitotic divisions in the presence of proper nutrients (*26*). In human female meiosis, reverse segregation often leads to euploid oocyte and PB2, suggesting normal MII segregation; the authors suggest that unresolved recombination intermediates may have both caused the reverse segregation in MI and facilitated proper MII segregation by linking the otherwise unrelated homolog chromatids (*7*). Given the 2% sequence divergence between B6 and Spret, it is possible that Mlh1 is limiting due to intensive mismatch repair and there may not be enough Mlh1 for resolving Holliday junction intermediates. However, we would like to emphasize that if recombined homolog chromatids co-segregate, they do not lead to LOH (**Fig. S1C**). Therefore, M2 diploids with LOH and mitotic segregation cannot be explained by co-segregation of unresolved intermediates.

We summarize the model in **Fig. 5D**. Note that of the 1,663 haploid cells we observed, the number of chromosomes with and vs. crossovers is half and half, indicating that they predominantly derive from the first three patterns of diploids and that diploids without interhomolog crossovers (the latter two patterns) do not contribute significantly to mature haploid cells that successfully complete MII. Also, we unexpectedly observe that mitochondrial copy number falls into a bimodal distribution in haploid cells (**Fig. 3H**), with one peak similar in copy number to M2 diploids that segregated meiotically (**Fig. S5E**), and the other similar in copy number to M2 diploids that segregated mitotically or other diploids that segregated mitotically and possibly with unobserved crossovers (**Fig. S1C**). At this time, we cannot rule out that this is merely a correlation without real biological meaning or due to physical characteristics of how easily the mitochondria dissociate from the bulk nuclei in different cell states.

Crossover hotness is a continuum and shaped by many factors. Consistent with previous findings, we observe that crossovers are hot in the middle of the chromosome arm and suppressed at centromeric and telomeric regions, and furthermore positively associated with Spo11 break hotness, particularly for hotspots where both B6 and Spret reference genomes contain PRDM9 binding sites (*20, 27*). Note that this only accounts for the PRDM9 sites bound by PRDM9 protein, and it is likely that the Spret copy of PRDM9 binds different sites and creates new meiotic DSB hotspots, and those are not accounted for in the current model. Crossover also prefers GC-rich regions, which is previously shown in yeast meiosis (*28, 29*). We have also found many new features that may affect crossover hotness: 1) existing CNV gains (*21*) between B6 and Spret strains are hot for crossovers as well as gene bodies and pseudogenic transcripts; 2) repeats behave differently: unlike in yeast, where rDNA is extremely cold for meiotic breaks (*30*), mouse rDNA is not suppressed for crossover; while DNA transposons, LTR, SINE and segmental duplications are hot for crossover, LINE and low complexity DNA and simple repeats are cold; 3) From ChIP-seq data, Dmrt6 (*31*) is an important gene regulating switch from mitotic to meiotic divisions of male germ cells and its binding sites are strongly positively associated with meiotic crossover hotness. TOP2B binding sites mapped in MEFs (*32*), however, are negatively associated; 4) In terms of higher order chromatin organization, CTCF binding sites (*33*) and replication domains mapped by Repli-seq (*34*) also positively correlate with hotness; 5) We found histone modifications, such as histone lysine pan-acetylation (Kac) and crotonylation (Kcro) (*35*), as well as H3K27me3 (*36*) to be strongly associated. However, Kac and Kcro positively correlate with one another while exhibiting opposite directions in the linear model, and could be counteracting each other in the model without being truly associated with crossover hotness. A few other well established marks such as H3K4me3, H3K27ac and H3K36me3 (*37, 38*) are not statistically significant in our model. It is possible that their effects have been represented by Spo11 break sites and are thus not significant given that Spo11 is already in the model. We emphasize that although the various features behave largely consistently between modeling approaches, we cannot assign any causality without further experiments.

In summary, sci-LIANTI expands the toolset for single-cell DNA sequencing, providing an effective balance between throughput and coverage. In this study, we have shown how sci-LIANTI can provide a systematic and quantitative view of meiotic recombination, with an unprecedented combination of throughput and resolution. We anticipate that sci-LIANTI can also be applied in other contexts where single cell genome sequencing is proving transformative, *e.g*. for dissecting the genetic heterogeneity and evolution of cancers.

## Acknowledgments

The raw data will be deposited with the Sequence Read Archive (www.ncbi.nlm.nih.gov/sra) and processed data will be deposited with Gene Expression Omnibus (www.ncbi.nlm.nih.gov/geo). We thank Dr. Giancarlo Bonora, Dr. Nancy Kleckner and members of the Shendure lab for helpful discussions. We thank Dr. Chongyi Chen, Dr. Dong Xing, Junyue Cao and Dr. Malte Spielmann for helpful technique suggestions, Anh Leith for her exceptional assistance in flow sorting, and Tom Reh’s lab for sharing the NIH/3T3 cell line. F.J.S. declares competing financial interests in the form of stock ownership and paid employment by Illumina. One or more embodiments of one or more patents and patent applications filed by Illumina may encompass the methods, reagents, and data disclosed in this manuscript. This work was funded by grants from the NIH (DP1HG007811 and R01HG006283 to J.S., DK107979 to J.S., W.S.N. and C.M.D., and GM046883 to C.M.D), and the Paul G. Allen Frontiers Group (Allen Discovery Center grant to J.S.). Y.Y. is a Damon Runyon Fellow supported by the Damon Runyon Cancer Research Foundation (DRG-2248-16). J.S. is an investigator of the Howard Hughes Medical Institute.

## Supplementary Materials

Supplemental Text

Figures S1-S9

Tables S1-S11

sci_lianti_inst.tar.gz

